# Mitochondria – insulin granule crosstalk controls the early stages of granule maturation

**DOI:** 10.64898/2026.02.23.707428

**Authors:** Styliani Panagiotou, Kousik Mandal, Sofia Amini, Kia Wee Tan, Samuel B. Stephens, Olof Idevall-Hagren

## Abstract

Insulin secretion from pancreatic β-cells is a tightly controlled process where hormone synthesis, granule formation and release are regulated in order to maintain whole body glucose homeostasis. Failure to produce or release insulin results in hyperglycemia that may develop into diabetes. Insulin-containing granules exist in different pools that have different propensity for release, yet what determined the fate of a granule after initial formation is not clear. In this study we aimed to identify key steps in the early life of an insulin granule that directs it towards release. Using two different methods for time-dependent labeling, we found that insulin granules shortly after budding from the trans-Golgi network associate with mitochondria. This organelle interaction involves the voltage-dependent anion channel (VDAC) and the vesicular nucleotide transporter (VNUT). Reduced VNUT expression prevented the recruitment of VDAC to insulin granules and redirected granules towards autophagy-dependent lysosomal degradation, resulting in reduced insulin content and impaired insulin secretion. These results show the requirement of granule-mitochondria crosstalk for normal progression through the early stages of the secretory pathway.

## INTRODUCTION

The islets of Langerhans are discrete clusters of endocrine cells that are scattered throughout the pancreas. β-cells constitute 60-80% of the islet cell population and synthesize and secrete insulin, which is the major blood glucose-lowering hormone that acts by promoting glucose uptake in peripheral tissues ^1^. Insulin secretion is a tightly regulated multi-step process that includes insulin granule (IG) biogenesis, maturation, and exocytosis ^2^. Insulin biosynthesis takes place in the endoplasmic reticulum (ER) where preproinsulin is converted to proinsulin, which is further modified in the Golgi apparatus and eventually packaged into immature IGs that bud from the trans-Golgi network ^3,4^. The acidification of the lumen of immature IGs activates prohormone convertases which catalyze the conversion of proinsulin into insulin, leading to IG maturation ^5^. Mature IGs are stored in the cytoplasm and consist of two distinct pools; the readily releasable pool which undergoes exocytosis upon stimulation and the reserve pool, which is located deeper in the cytosol ^6^. Three main factors are considered to increase the propensity for IG exocytosis; the composition of the IG membrane, the localization inside the cell and the IG age. Release-competent insulin granules are required to have a specific lipid and protein profile which facilitate priming, docking, and fusion with the plasma membrane upon a triggering increase in the intracellular Ca^2+^ concentration ^7,8^ . While most protein sorting and lipid modifications occurs in conjunction with granule budding from the trans-Golgi network, the granule composition also continuously changes as it moves along the secretory pathway ^5^. Early steps determine whether the granule is directed towards release or storage, and the former are then transported along microtubules to the plasma membrane where they are released shortly after synthesis ^9–13^. Recent studies have revealed age-specific proteomic and lipidomic IG characteristics that favor release. For example, negatively charged lipids on newly synthesized granules allows the binding of effectors, such as KIF5b, that increase their competence for microtubule-mediated transport ^8^. Lipid modification of the granule membrane has been proposed to depend both on enzymatic reactions and non-vesicular transfer ^14,15^, although for the most part it is not known how these changes occur. The granule proteome also changes during maturation, both by active sorting from the granule surface and by recruitment via protein-protein or protein-lipid interactions^8^. We recently reported that IGs form physical contacts with the endoplasmic reticulum (ER), and that transport of cholesterol from the ER to IGs at these membrane contact sites was important for granule biogenesis and release ^15^. Other membrane contact sites, including those formed between ER and mitochondria, are essential for beta cell function by facilitating the exchange of lipids and Ca^2+^ ^16,17^. Recently, more complex contact sites involving three or more organelles have been described, including contacts between the ER, mitochondria and Golgi-derived transport vesicles ^18^. There are also both old and new observations of close proximities between mitochondria and IG, but the functional significance of these contacts is unclear, and it is not known if they involve a specific granule pool or play a role in granule maturation ^19,20^.

The rate of IG biogenesis likely exceeds that of exocytosis and mechanisms where IGs are degraded are therefore acting in parallel with biogenesis to maintain the size of the IG pools ^21^. Both macroautophagy or more specialized versions of this process contribute to IG turnover by promoting lysosomal degradation ^22,23^. In addition to this, proinsulin and insulin can be specifically removed from IGs by ELAPOR1, which acts as a sorting receptor for lysosomal degradation ^24^. Disturbances in these degradative pathways have been observed in diabetes ^22^, yet it remains largely unknown how IG turnover is regulated in normal physiology and how beta cells are able to balance IG biogenesis and degradation.

In this study we show that IGs encounter mitochondria shortly after budding from the Golgi, likely stabilized via an interaction between VNUT on the IG membrane and VDAC on the mitochondrial membrane. Loss of VNUT results in reduced granular ATP content, impaired IG exocytosis and reduced insulin secretion, largely due to dramatic loss of insulin as a consequence of enhanced autophagy and lysosomal degradation.

## MATERIALS AND METHODS

### Reagents and plasmids

NPY-mNG, VAMP2-pHluorin and GFP-Rab3a were gifts from Sebastian Barg (Uppsala University, Sweden), mCherry-tandem C2-domains of PKC (C1aC1b) and mCherry-GRP1-PH was previously described ^25,26^. proCpepRUSH system was a gift from Samuel Stephens (University of Iowa, Iowa, USA) (Boyer et al., 2023). R-GECO was a kind gift from Robert Campbell ^27^. mApple-TOM20 was a kind gift from Michael Davidson (Addgene plasmid #54955). NPY-Halo was created by removing mNeonGreen (mNG) from NPY-mNG and replacing it with Halo from a C-terminal Halo-tag (pHalo-N1) plasmid using AgeI/NotI. NPY- Halo was inserted in pENTR20 vector using NheI/NotI restriction sites. This vector was recombined using Gateway LR reaction (Invitrogen) into a customized Gateway-enabled pCDH vector (System Biosciences) and expressed under control of a EF1α promoter. NPY-mCh-mNG was created by PCR and Gibson Assembly of an mCherry insert and a 20 amino acid rigid spacer into the AgeI site of NPY-mNeonGreen. The following antibodies and dyes were used in the study: Rab3 monoclonal (catalog no. 107 111, Synaptic Systems, host: mouse, 1:500), TOM20 polyclonal (catalog no. 11802-1-AP, Proteintech, host: Rabbit, 1:400), insulin polyclonal (catalog no. A0564, Dako, host: guinea pig, 1:1000), VDAC (catalog no. MA533205, host: rabbit, 1:200, Invitrogen), VDAC1 (catalog no. 66345-1-lg, Proteintech, host: mouse, 1:200), VDAC2 (catalog no. 66388-1-lg, Proteintech, host: mouse, 1:200), VNUT polyclonal (catalog no. ABN83, Merck, host: guinea pig, 1:200). SiR lysosome dye (catalogue no. SC012, Spirochrome). Where eGFP-tagged proteins were visualized in fixed samples, the secondary antibody incubation included ChromoTek GFP-Booster Alexa Fluor 488 (catalog no. gb2AF488, Proteintech, nanobody, 1:500). Cal590 AM was from AAT Bioquest (catalog no. 20511). SiR-lysosome was from Spirochrome. All salts were from Sigma Aldrich. The following chemical were used: SAR405 (catalogue no. 5.33063, Sigma-Aldrich), clodronate (catalog no. 233183- 10MG, Sigma-Aldrich), choloquine (Sigma-Alrich, catalog no. 233183), JFX549 and JFX650 as Halo-tag ligands (kind gifts from Luke Lavis ^28^).

### Cell culture

The mouse β-cell line MIN6 (passages 18–30; gift from Jun-ichi Miyazaki, Kumomoto University, Japan) ^29^ was cultured in the Dulbecco’s modified Eagle’s medium (DMEM) supplemented with 4.5 g/L glucose, 2mM L-glutamine, 100U/mL Penicillin, 100μg/mL Streptomycin, 50μM 2-mercaptoethanol, and 15% fetal bovine serum (complete culture medium) (all the above from Life Technologies). The cells were maintained in a humidified atmosphere at 37°C and 5% CO_2_.

### Transfection and siRNA-mediated knockdown

Reverse transfections were performed by combining 50 μL of OptiMEM containing 0.1-0.2 μg DNA and 0.25-0.5 μL Lipofectamine 2000 (all from Life Technologies) with 50 μL OptiMEM containing 200,000 cells. The mix of transfection reagents, plasmids and cells was seeded onto the center of a 25 mm glass coverslip. The transfection was terminated after 5 h using 2-4 mL of complete culture medium, and imaging was done between 22 and 30 h after transfection. Small interfering RNA (siRNA) knockdown experiments were performed by double transfection reaction. MIN6 cells were transfected in 6 well-plates with siRNA using Lipofectamine 2000, followed by a second round of transfection 3-4 h later using RNAiMax (Life Technologies) according to the manufacturer’s instructions with final siRNA concentration of 20 nM. VNUT knockdown was performed with siRNA targeting mouse gene sequence (Ambion Silencer Select Pre-Designed siRNAs; Cat#:4390771, s106087). The sequences that were used are sense (5’→3’): CAACCACAGUGGUAUUUCAtt and antisense (5’→3’): UGAAAUACCACUGUGGUUGaa. VDAC knockdown was performed with siRNA targeting mouse gene sequence (Ambion Silencer Select Pre-Designed siRNAs; Cat#:4390771, s75922 and s75920 for VDAC1 and s75924 and s75925 for VDAC2). For the control experiments, the cells were transfected with Silencer Select Negative Control #1 siRNA; Cat no: 4390843. Cells destined for imaging were plated on glass coverslips while cells designated for RT-qPCR were plated on poly-L-lysine-coated 6-well plates. The transfection was stopped by adding complete culture medium.

### Pancreatic islet isolation and culture

All animal experimental procedures were approved by the local ethics committee for use of laboratory animals in Uppsala, Sweden. The pancreata were collected from 4 to 8-month-old C57Bl6J mice and digested using collagenase P digestion on a shaker at 37°C. Individual islets of Langerhans were hand-picked under a stereo microscope. After isolation, the islets were cultured for 1–2 days in RPMI 1640 medium (Gibco) with 5.5 mM glucose supplemented with 100 U/ml penicillin, 100 μg/ml streptomycin and 10% FBS at 37°C in a humidified atmosphere containing 5% CO2. The Nordic Network for Clinical Islet Transplantation at the Academic hospital in Uppsala provided the isolated human islets from normoglycemic cadaveric organ donors. The human islets were picked by hand under a stereomicroscope and cultured in CMRL Medium (Gibco) with 5.5 mM glucose supplemented with 10% FBS, 100 U/ml penicillin and 100 μg/ml streptomycin at 37°C and containing 5% CO_2_ for up to one week. All experiments were approved by the Uppsala human ethics committee.

### Viral transduction of MIN6 cells and mouse pancreatic islets

MIN6 cells and mouse pancreatic islets were infected with ∼2–10 × 10^7^ IFU/mL adenovirus for 3 h 8at 37°C in a humidified atmosphere containing 5% CO_2_. After 3 h, glass coverslips with seeded MIN6 and sterile dishes containing free-floating pancreatic islets were washed with medium and cultured for at least 48 h to allow the expression of proCpepRUSH or NPY-mCherry-mNeonGreen.

### Immunofluorescence

Cell monolayer, grown on coverslips, and pancreatic islets were fixed in 4% paraformaldehyde in PBS for 20 min at RT, followed by three 5-min washes using PBS. Samples were permeabilized using 0.1% Triton X-100 in PBS for 5 mins, followed by 3 washes of buffer. After samples were blocked for 1hr in buffer containing 2% BSA in PBS, primary antibody incubation was performed overnight at 4°C in blocking buffer. Unbound primary antibody was removed during the three 5-min washes of buffer and samples were incubated for 1h with secondary antibody in blocking buffer. Removal of excess secondary antibody was carried out by three washes of buffer and samples were mounted in Prolong Glass (Invitrogen) and left to dry for at least 20hr before imaging.

### Quantitative RT-PCR

Knockdown efficiency was determined by quantitative RT-PCR using SYBR Fast SYBR Green Master Mix (Fisher Scientific) and the following primers: VNUT-fwd: 5ʹAAGGAGGCTGGTATCGTGC-3ʹ, VNUT-rev: 5-TGGTCAGGGCTGGAAAGTAG-3’; GAPDH-fwd: 5ʹ- ACTCCACTCACGGCAAATTC-3’; VDAC1-fwd: 5’- TCACCGCCTCCGAGAACAT-3ʹ; VDAC1-rev: 5’-ACGTCAGCCCATACTCAGTC-3ʹ; VDAC2-fwd: 5’- ACTCTGAGGCCTGGTGTGAA-3ʹ; VDAC2-rev: 5’- AGCCTCCAATTCCAAGGCAA-3ʹ; GAPDH-rev: 5ʹ-TCTCCATGGTGGTGAAGACA-3’. PCR reactions were performed using QuantStudio 5 RT-PCR System (Applied Biosystems). Results are expressed as ΔΔCt, normalized to the expression in control samples.

### Treatment with inhibitors

The following inhibitors were used in this study: Clodronate (120 nM; catalog no. 233183- 10MG, Sigma-Aldrich), Chloroquine (20 μM; catalog no. C6628, Sigma-Aldrich) and VPS34 Inhibitor, SAR405 (10 μM; catalogue no. 5.33063, Sigma-Aldrich). All inhibitors were added directly to culture medium and cell were treated for 12-16 h unless otherwise specified.

### Proximity ligation assay

PLA was performed according to the manufacturer’s recommendation (Navinci NaveniFlex MR). Primary antibodies against VDAC (catalog no. MA533205, host: rabbit, 1:200, Invitrogen), Rab3 (catalog no. 107 111, mouse monoclonal, Synaptic Systems) and VNUT (catalog no. ABN83, Merck, host: guinea pig) were used at 1:200 dilution.

### Pulse-chase experiments

In samples expressing NPY-Halo, pre-existing insulin granules were labelled with 400 nM of Halo-tag ligand JFX650 dissolved in culture medium for 20 min at 37°C (1st pulse) ^28^. The unbound ligand was removed during two 5-min washing steps in culture medium. New NPY- Halo positive insulin granules that were synthesized withing the next 1h (1st chase) were subsequently labeled with 400 nM of Halo-tag ligand JFX549 (2nd pulse) and the excess ligand was washed as previously described. Unless it is stated otherwise, the samples were fixed 30 min after the 2nd pulse. During this time the newly-synthesized granules bud from the Golgi apparatus but remain in the proximal region.

### Insulin content measurement

Total insulin from the MIN6 cells were extracted using 0.18M acid ethanol. Total insulin content was measured using mouse insulin ELISA (catalogue no. 10-1247-01, Mercodia) and normalized to the cell number for each sample, determined by counting of viable cells using a Countess 3 automated cell counter (Invitrogen).

### Confocal microscopy

The immunolabeled samples were imaged using both a spinning-disk confocal microscope for live-cell imaging and laser-scanning confocal microscope for fixed samples. The first one is a Nikon Eclipse Ti-2 equipped with a Yokogawa CSU-10 spinning disc confocal unit and a 100×/1.49-NA plan Apochromat objective (Nikon). Excitation light was provided by 491 and 561-nm diode-pumped solid state (DPSS) lasers and a 640-nm diode laser (all from Cobolt/Hübner Photonics). For the selection of the excited light, electronic shutters were used (SmartShutter, Sutter Instruments) and the emitted fluorescence light was separated with the following filters placed in a filter wheel controlled by a Lambda 10-3 unit (Sutter Instruments); 530/50-nm (GFP, mNG, Alexa 488), 590/20-nm (mCherry, Alexa 568, JFX549) or 650LP filter (JF650, Alexa 647). Images were captured using a back-illuminated electron-multiplying charge-coupled device (EMCCD) camera (DU-888; Andor technology) controlled by MetaFluor software (Molecular Devices). Fixed samples were imaged using an upright laser scanning confocal microscope (Zeiss LSM700) equipped with a 63x/ 1.4 oil-NA objective, 405 (Hoechst)/488 (GFP-booster, Alexa 488), 555 (Alexa 561), 633 (Alexa 647) lasers and two Photomultiplier tubes (PMTs). Raw data were acquired with ZEN (black edition) software (BioVis facility, Uppsala University, Sweden).

### TIRF microscopy

All experiments, unless otherwise stated, were performed at 37°C in an experimental buffer consisting of 125 mM NaCl, 4.9 mM KCl, 1.2 mM MgCl_2_, 1.3 mM CaCl_2_, 25 mM HEPES, 3 mM D-Glucose, and 0.1% BSA (pH 7.40). Cells were preincubated in the experimental buffer for 30 min before experiments, and continuously perfused with the same buffer at the stage of the microscope where the temperature was maintained by a custom-made stage heater and an objective heater. TIRF imaging was performed on an inverted Nikon Ti-E equipped with a TIRF illuminator and a 60×1.45-NA objective (all Nikon). Diode-pumped solid-state lasers provide the excitation light for pHluorin (491-nm) and mCherry (561-nm). After they get merged with dichroic mirrors (Chroma technologies), excitation light separation is conducted via bandpass filters placed in a filter wheel (Sutter Instruments Lambda 10-3) and light is directed into a fiber optic cable and delivered to the TIRF illuminator. Excitation light is separated through the objective with a dichroic mirror (ZET405/488/561/640m-TRFv2, Chroma Technologies) and emission light was separated using interference (530/50 nm, 590/20-nm) filters (Semrock) mounted in a filter wheel (Sutter instruments Lambda 10-3). An electronic shutter was used to control for the beam blockage during image acquisition. Emission light was collected using an Orca-AR camera controlled by MetaFluor software (Molecular Devices Corp.). TIRF microscopy recordings of VAMP2-pHluorin and mCherry-GRP1-PH fluorescence (Fig. 4) was performed on a custom-built prism-based setup or an objective-based system as previously described ^30^. Diode-pumped solid-state lasers (Cobolt, Solna, Sweden) provided excitation light for VAMP2-pHluorin (491 nm) and mCherry-GRP1-PH (561 nm). Emission wavelengths were selected with filters (542/25 nm for GFP and 597/20 for mCherry; Semrock, Rochester, NY) mounted in a filter wheel (Sutter Instruments). Fluorescence was detected with a back-illuminated EMCCD camera (DU-897, Andor Technology) under MetaFluor (Molecular Devices Corp, Downington, PA) software control. Images or image pairs were acquired every 5 seconds.

### Image analysis

The acquired confocal and TIRF images were analyzed using ImageJ (Fiji) and Cell Profiler. For analysis of VAMP2-pHluorin and mCherry-C1aC1b, regions of interest were assigned based on the cell footprint. Changes in the intensity of the fluorescence signal were exported to Excel. All values were background corrected and normalized to pre-stimulatory level (F/F0). For statistical comparisons, ΔF was calculated as the peak fluorescence divided by the pre- stimulatory fluorescence and compared between conditions. The enrichment of various fluorescently tagged proteins on insulin granules was calculated using a pipeline built in Cell Profiler. Old and new insulin granules were segmented using the proper channel and granules inside cells negative for the protein of interest were discarded. The median fluorescence of the protein of interest was measured in a 2-pixel wide ring around the insulin granules, corresponding to the background. The mean fluorescence of the protein in the insulin granule region divided by its background allowed for the calculation of the enrichment of the protein of interest on the individual insulin granules. A threshold of 2 standard deviation (SD) from the mean was set for the enrichment in the positive insulin granules. When cells were treated, the SD of the basal condition were computed and applied to the corresponding treatment conditions. To measure co-localization of the fluorescent signal of a protein between two channels, the Just Another Co-localization plugin (JACoP) in ImageJ was applied to generate the Mander’s coefficient values. For the lysosomal degradation experiments, *mCherry only area* (mCherry+ in the figure) was quantified by thresholding both mNG and mCherry channels at mean + 2SDs for each individual cell. Binary masks were generated for both channels and mNG-positive area was subtracted from the mCherry-positive area to obtain the *mCherry-only area*. *mCherry only area* was normalized to the total cell area to account for cell size variability.

Average intensity within the *mCherry only area* was normalized with the median of control sample for individual replicates.

### Quantification and statistical analysis

The statistical analysis in this study was performed using GraphPad prism software (version 9.5.1). First the distribution of the raw data was assessed using the Kolmogorov-Smirnov test. For non-normally distributed data, non-parametric Mann-Whitney signed-ranked test was performed while for multiple comparisons Friedman and Kruskal-Wallis test were applied followed by Dunn–Šidák’s multiple comparisons post hoc test. For all analyses, at least three independent experiments (biological replicates) were performed on three separate days.

## RESULTS

### Time-dependent interactions between mitochondria and newly synthesized insulin granules

Newly synthesized insulin granules (IGs) undergo extensive remodeling of both protein and lipid content as part of the maturation process. Such remodeling is energy demanding and recent work has shown that there is a structural basis for direct interactions between IGs and ATP-producing mitochondria ^19^. To determine the dynamics of this interaction we expressed the IG marker NPY-mNeonGreen and the mitochondria marker Tom20-mApple in MIN6 beta cells and followed them over time by confocal microscopy. We found that 40% the IGs in the cell center were in close proximity to mitochondria, while less then 20% of the docked IG at the cell periphery exhibited this feature (Fig. 1A-C). The more centrally located IGs may represent granules that were recently synthesized or granules belonging to the reserve pool. To determine if the interactions between mitochondria and IGs depend on granule age, we performed pulse-chase experiments in MIN6 β-cells expressing NPY-Halo. First, we labelled all pre-existing granules with JFX650 and then chased the cells for 60 min in the absence of dye followed by a second round of labeling using the spectrally separable fluorophore JFX549 (Fig. 1D; Fig. S1A, C). The cells were then chased for 30 min, fixed, immunostained against the mitochondrial protein Tom20 and imaged with confocal microscopy. We found both old and new IGs in close proximity to mitochondria, but the fraction was twice as high for IGs of younger age (Fig. 1D,E). Similar to the observation in live cells, the interactions between

**Figure 1.**
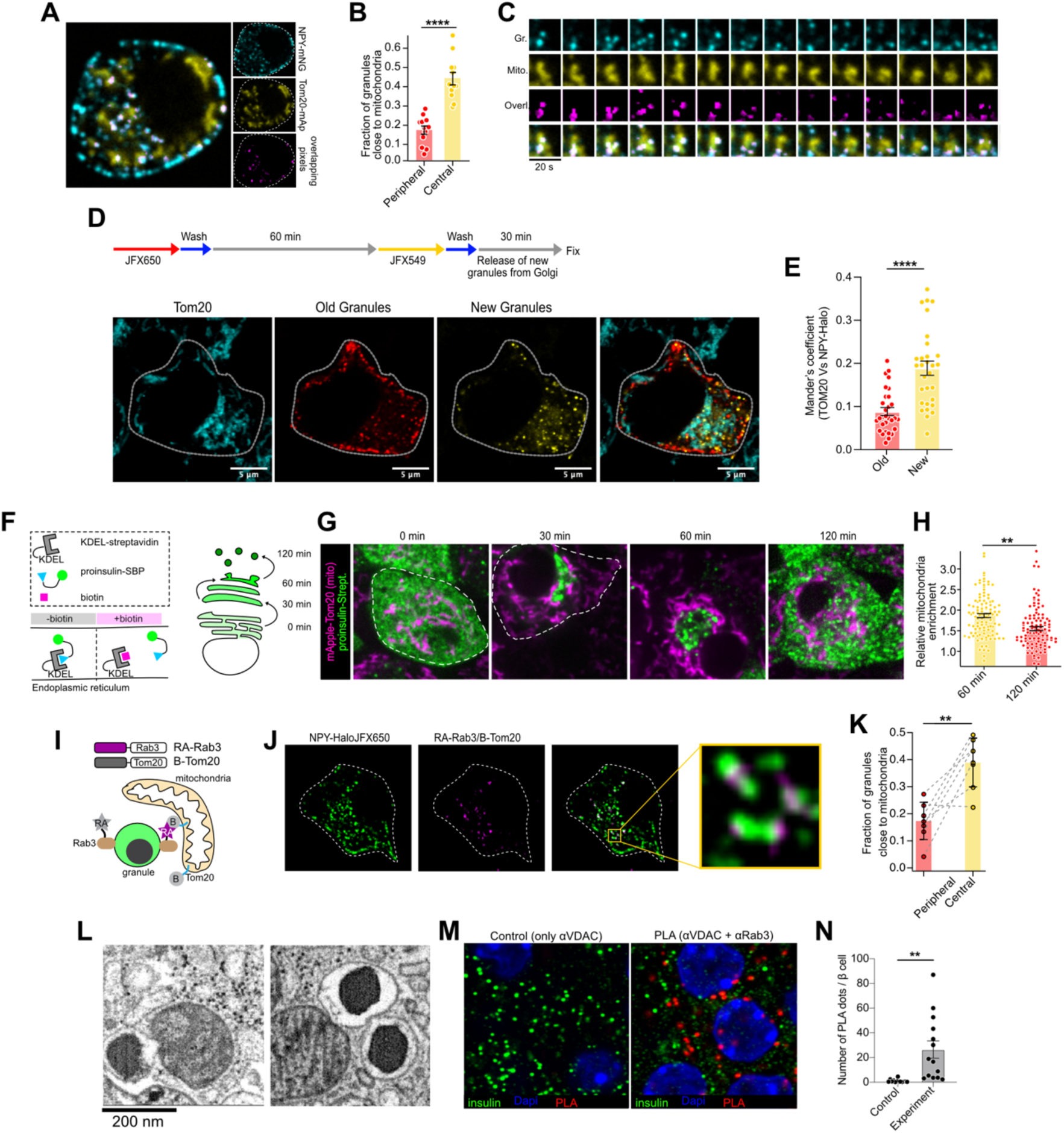
Interactions between newly synthesized granules and mitochondria (A) Confocal microscopy image of a MIN6 cell expressing NPY-mNeonGreen (cyan) and Tom20-mApple (yellow). Pixel overlaps between cyan and yellow is shown in magenta.(B) Fraction of NPY-mNeonGreen-containing granules in close proximity to mitochondria (Tom20-mApple) (n=12 cells from three experiments; means ± SEM; Student’s paired t-test; ****P<0.0001).(C) Montage of images from a live cell showing the spatio-temporal relationship between granules (cyan) and mitochondria (yellow). Pictures in magenta show pixel overlap between cyan and yellow. (D) Workflow for pulse-chase labeling of insulin granules based on age (top) and confocal microscopy images of a MIN6 cells expressing NPY-Halo and labeled according to the workflow followed by immunostaining against the mitochondrial marker Tom20 (cyan). Scale bars: 5 μm. (E) Mander’s coefficients denoting the fraction of Tom20 overlapping with new (JFX549- labeled) and old (JFX650-labeled) insulin granules (means ± SEM, ****P<0.001; Kruskal–Wallis test; n = 30 cells). (F) Cartoon showing the principle of the RUSH system. (G) Confocal microscopy images of MIN6 cells expressing spGFP-proCpepRUSH (green) and the mitochondrial marker Tom20-mApple (magenta). Images are from 0, 30, 60 and 120 min post-biotin addition (200 μM). (H) Relative mitochondria enrichment on spGFP-proCpep-positive granules 60 and 120 min after biotin addition (n=111 cells; three experiments; means ± SEM; Student’s unpaired t-test; ** P<0.01). (I) Cartoon showing the principle behind detection of granule-mitochondria contacts in live cells using the split-fluorophore RA-GB system. (J) Confocal microscopy images of a MIN6 cell expressing the granule marker NPY- Halo^JFX650^ and the granule-mitochondria proximity detector (magenta). Boxed area is magnified to the left and shows how mitochondria-granule proximities are restricted to a small surface on the granule. (K) Fraction of granules in proximity to mitochondria based on their subcellular localization (peripheral or central) (n=8 cells; three experiments; means ± SEM; Student’s paired t-test; **P<0.01). (L) FIB-SEM images showing the proximity between insulin granules and mitochondria in a mouse β-cell. Scale bar: 200 nm. (M) Confocal microscopy images of mouse islets with insulin immunostaining shown in green and mitochondria-granule proximity sites detected with PLA, using antibodies against the mitochondria (VDAC) and insulin granules (Rab3a), shown in red. In control experiments, only the VDAC antibody was used. Scale bar: 50 μm. (N) Average number of PLA puncta per insulin-positive cell (single confocal plane; means ± SEM, n = 14, >30 islets for each experiment, unpaired, two-tailed Student’s t test, **p < 0.01).

mitochondria and newly synthesized IGs occurred primarily in a perinuclear region; a location that resembles that of the Golgi. To more accurately monitor the formation of IGs and their relationship to mitochondria, we expressed a proinsulin trafficking reporter system based on the retention using selective hooks (RUSH) methodology ^31,32^ (Fig. 1F). Briefly, sfGFP-tagged proinsulin was fused to a streptavidin-binding protein (SBP) and coexpressed with streptavidin fused to an ER-retention sequence (KDEL). Under resting conditions, the interaction between streptavidin and SBP prevents export of proinsulin from the ER. Upon addition of biotin, this interaction is reversed, leading to synchronized trafficking of proinsulin. 60 min after the addition of biotin, fluorescent puncta appeared in the perinuclear region, consistent with export from the Golgi apparatus (Fig. 1G and Fig. S1B). Co-expression of mApple-Tom20 further revealed close apposition between mitochondria and newly synthesized IGs at the trans-Golgi network; a proximity that was reduced over time as the IGs matured (Fig. 1G,H). To directly visualize interactions between mitochondria and IGs in living cells, we adopted a split-fluorophore methodology where two non-fluorescent components were anchored to the mitochondria and IG surface, using Tom20 and Rab3 respectively, that, when in proximity to each other, refold into a red-light emitting protein (Fig. 1I) ^15,33^. Importantly, Rab3 was present on both old and new insulin granules, consistent with previous observations ^15^ (Fig. S1D). Co-expression with the IG marker NPY-Halo^JFX650^ demonstrated the presence of mitochondria in close proximity to IGs in living cells, and showed that these interactions primarily involved IGs in the cell center (Fig. 1J,K). Furthermore, examination of FIB-SEM data sets (openorganelle.janelia.org) ^34^ confirmed the existence of close contacts between IGs and mitochondria in mouse islet beta cells (Fig. 1L). Contacts between mitochondria and other organelles often involve mitochondrial VDAC ^35^, and we therefore tested if this was also the case for IG-mitochondria contacts in mouse islets using proximity ligation assay and antibodies against VDAC (mitochondria) and Rab3 (IG). Confocal microscopy imaging revealed around 20 fluorescent puncta per imaging plane and cell, while less than 1 puncta was seen in control cells where only one of the antibodies was included (Fig. 1M,N and Fig. S1E-F). Together, these results show that there are close proximities between IGs and mitochondria that preferentially involves newly synthesized granules.

### VNUT and VDAC colocalize at newly synthesized insulin granules

The close proximity between IGs and mitochondria early in the secretory pathway indicate that this interaction is important for granule maturation. The ATP concentration in secretory granules is around 100 times higher than in the cytosol, and uptake occurs through the vesicular nucleotide transporter (VNUT; Slc17A9)^36^. Since mitochondria are the main source of ATP, we next investigated a scenario where IG become adjacent to mitochondria to facilitate ATP uptake. Confocal microscopy images of MIN6 cells showed the presence of VNUT on most, but not all, IGs. (Fig. 2A), and IGs positive for VNUT were often in proximity to mitochondria, identified by VDAC1 immunolabeling (Fig. 2B). Moreover, pulse-chase labeling of MIN6 cells expressing NPY-Halo showed that the overlap between VDAC and insulin was highest for granules that recently departed the Golgi (Fig. 2D), consistent with the observation that mitochondria preferentially interact with newly synthesized IGs (see Fig. 1). Next, we used the RUSH methodology to synchronously release proinsulin into the secretory pathway, fixed the cells at 30, 60 and 120 min post-biotin addition and immunostained against VDAC and VNUT. Confocal microscopy imaging showed that VNUT colocalized with proinsulin after 30 min, and after 60 min both VDAC and VNUT colocalized with proinsulin. After 120 min, proinsulin- containing granules positive for VNUT had formed, but VDAC maintained a perinuclear distribution and did not colocalize with VNUT or proinsulin (Fig. 2E). These results are consistent with the existence of IG-mitochondria contacts early in the secretory pathway, potentially at sites where VDAC and VNUT are positioned adjacent to each other. To experimentally test this, we performed proximity ligation assays on mouse islets using primary antibodies against VNUT and VDAC followed by immunostaining against insulin to identify β- cells. Around 30 fluorescent puncta per imaging plane and cell was visible using a VDAC1 antibody, but less than 5 puncta were seen when a VDAC2 antibody was used instead (Fig. 2F,G and Fig. S2A,B), indicating close proximities between granular VNUT and mitochondrial VDAC1.

**Figure 2.**
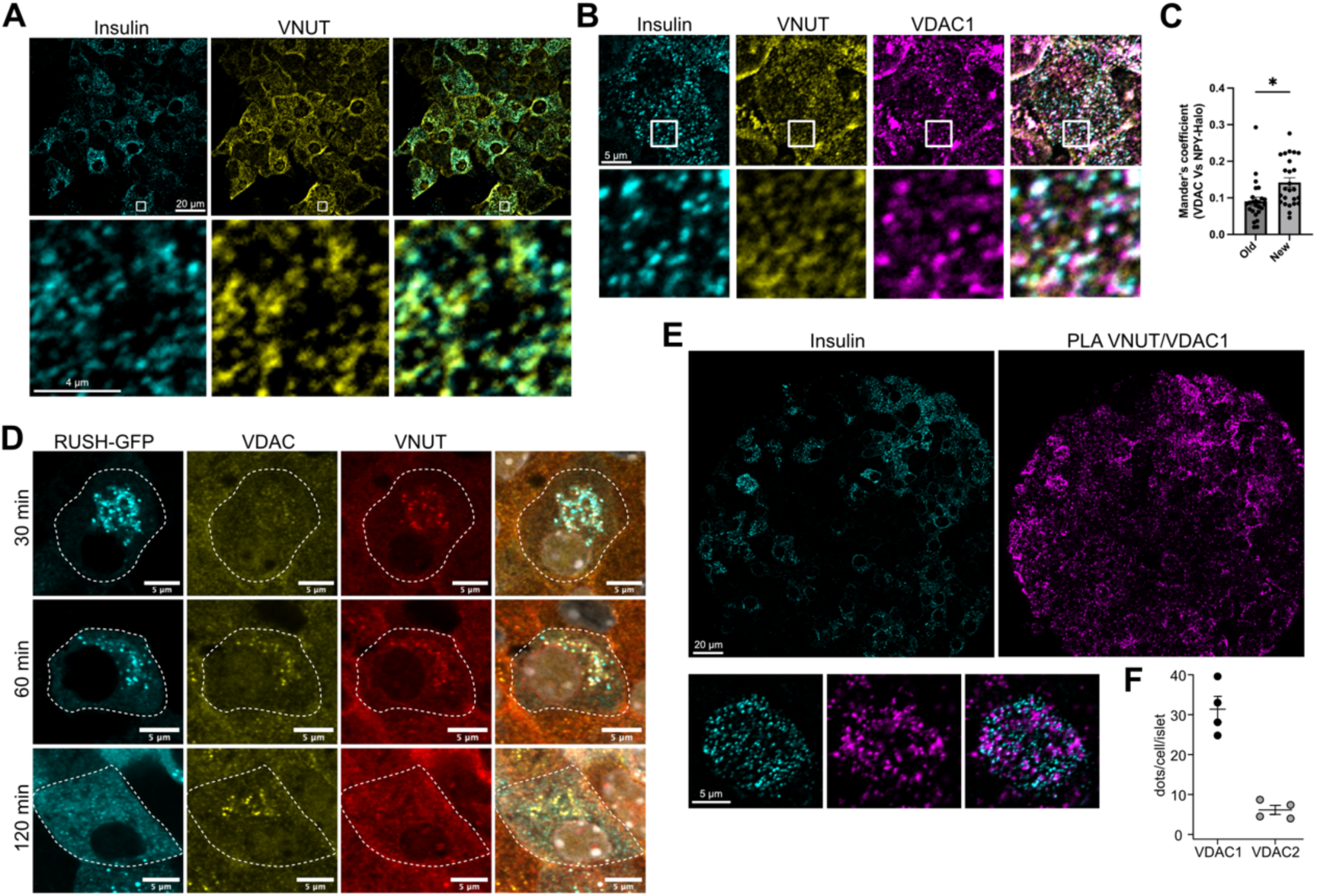
VNUT and VDAC associate with newly formed insulin granules (A) Confocal microscopy images of a MIN6 cell immunostained against insulin (green) and VNUT (magenta). (B) Confocal microscopy images of a MIN6 cell immunostained against insulin (green), VDAC (yellow) and VNUT (red). (C) Quantification of the colocalization between VDAC and NPY-Halo. A pulse-chase protocol for Halo labeling was employed where pre-existing granules were labeled with JFX549 (old) and newly synthesized granules with JFX650 (old) (means ± SEM; n = 25 cells from three experiments; Mann-Whitney U-test; *p < 0.05). (D) Confocal microscopy images of mouse islets expressing the AdRIP-proCpepRUSH 30, 60 and 120 min post-biotin addition (200 μM; cyan) and immunostained against VDAC (yellow) and VNUT (red). (E) Confocal microscopy images from a mouse islet showing PLA signal between VNUT and VDAC1 (magenta) and insulin immunoreactivity (cyan). The white region is expanded below. (F) Average number of PLA puncta (using antibodies against VNUT and VDAC1 or VNUT and VDAC2) per insulin-positive cell (single confocal plane; means ± SEM, n = 4 experiments).

### Reduced VNUT expression prevent VDAC localization to newly synthesized IG and leads to insulin deficiency

To investigate the putative role of VNUT in mitochondria-IG tethering, we reduced VNUT expression by siRNA-mediated knockdown (Fig. S2C-E). Expression of the proinsulin-trafficking reporter revealed, as expected, reduced VNUT immunoreactivity on newly formed IGs and reduced overlap between VNUT and VDAC in VNUT knockdown cells compared to control cells (Fig. 3A-D). Interestingly, the colocalization between mitochondrial VDAC and proinsulin seen in control cells (Fig. 2E and Fig. 3A) was completely abolished in VNUT knockdown cells (Fig. 3B,C). Using MIN6 cells expressing NPY-Halo and a pulse-chase protocol we found a reduction in the ratio of newly synthesized IGs to old IGs in VNUT KD cells (Fig. 3E,F), while the formation of new IGs at the trans-Golgi network, determined using the RUSH methodology, was unaffected (Fig. 3G). This indicates a role of VNUT in IG maturation or turnover but not in budding from the trans-Golgi network. In contrast, we observe impaired proinsulin trafficking in cells with reduced VDAC1 expression, indicating that VDAC1 has roles in IG formation beyond its interaction with VNUT (Fig. S2E,F). To further investigate the mechanism by which VNUT controls IG maturation, we immunostained control cells and cells with reduced VNUT expression against proinsulin and insulin. While proinsulin immunoreactivity was only slightly reduced in VNUT KD cells, insulin immunoreactivity was dramatically lower, resulting in reduced insulin:proinsulin ratio (Fig. 3I-K). This was due to both a reduction in the number of IGs and in the insulin content of the remaining granules, resulting in severe insulin deficiency (Fig. 3L-O). Next, we immunolabeled mitochondria in control and VNUT knockdown cells and visualized them with confocal microscopy. While the overall mitochondrial density was the same, the mitochondrial network complexity was reduced and mitochondrial branch length increased in VNUT knockdown cells (Fig. S3A-D). This was accompanied by slightly increased mitochondrial membrane potential determined using the voltage-sensitive dye TMRM following addition of the uncoupler FCCP (Fig. S3E-G). Together, these results show that VNUT is essential for maintaining insulin content in β-cells, but also indicate a role of VNUT in the regulation of mitochondrial function.

**Figure 3.**
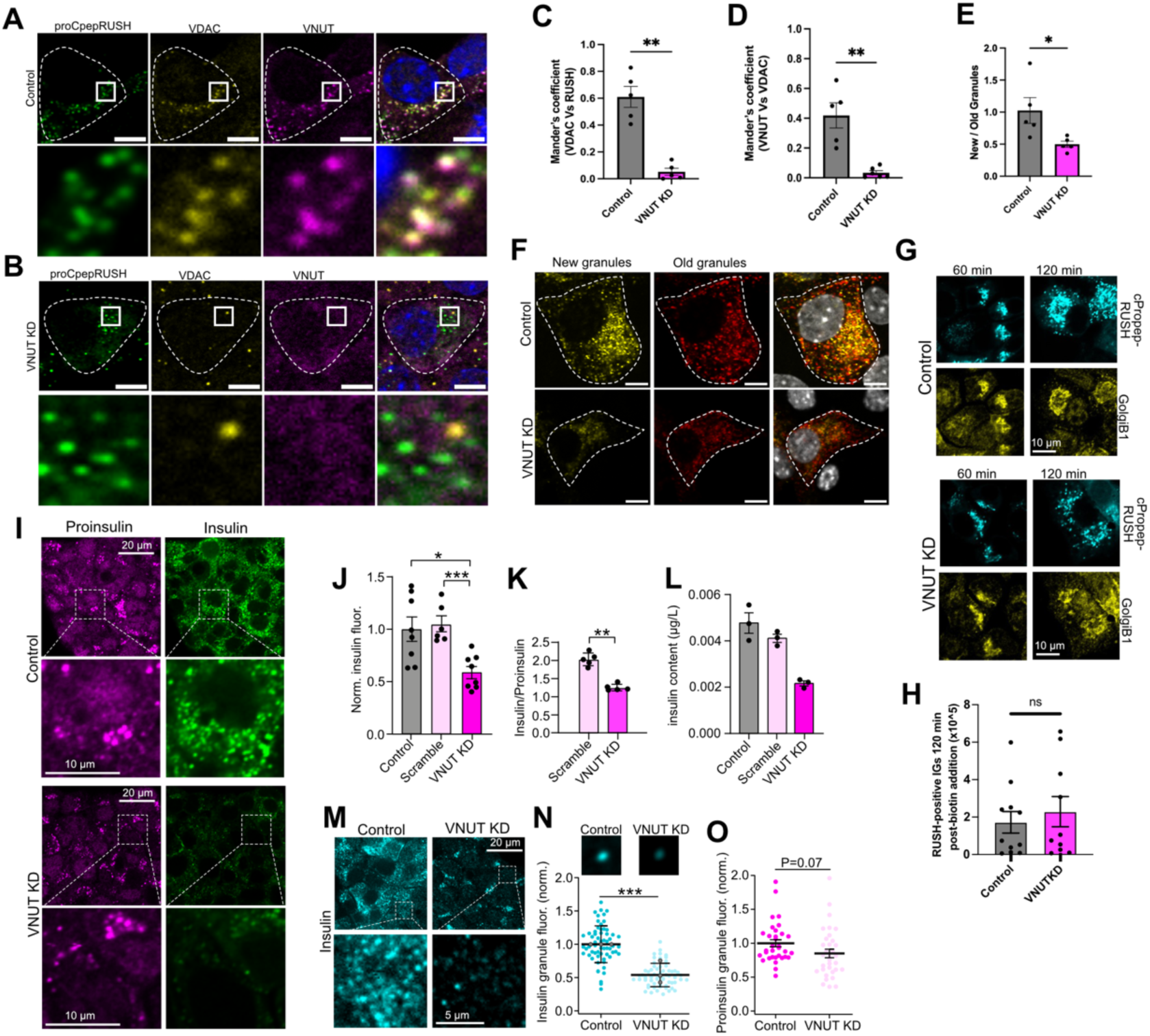
VNUT is required for IG-mitochondria interactions and IG maturation. (A) Confocal microscopy images of control MIN6 cells expressing AdRIP-proCpepRUSH 60 min post-biotin addition (green; 200 μM) and immunolabeled against VDAC (yellow) and VNUT (magenta). Enlarged sections are magnified from boxed regions. Scale bars: 5 μm. (B) Confocal microscopy images of VNUT KD MIN6 cells expressing AdRIP-proCpepRUSH 60 min post-biotin addition (green; 200 μM) and immunolabeled against VDAC (yellow) and VNUT (magenta). Enlarged sections are magnified from boxed regions. Scale bars: 5 μm. (C) Mander’s coefficient showing the overlap between VDAC and proCpepRUSH in control (grey) and VNUT KD cells (magenta) (means ± SEM; n=6 experiments, >50 cells, **P<0.01). (D) Mander’s coefficient showing the overlap between VNUT and VDAC in control (grey) and VNUT KD cells (magenta) (means ± SEM; n=5 experiments, >50 cells, **P<0.01). (E) Quantification of the ratio between the number of new and old insulin granules in control (grey bars) and VNUT KD (magenta bars) cells (means ± SEM; n=5 experiments, >50 cells). (F) Confocal microscopy images of MIN6 cells expressing NPY-halo labeled with JFX650 (old granules; red) and JFX549 (new granules; yellow) in control (upper panel) and VNUT KD (lower panel) cells. Scale bars: 5 μm. (G) Confocal microscopy images from control and VNUT KD cells expressing proCpepRUSH (cyan) that have been incubated with biotin for 60 and 120 min before fixation and immunostaining against GolgiB1 to visualize the Golgi. (H) Quantification of the number of proCpep-spGFP-positive granules 120 min after the addition of biotin in control and VNUT KD cells (means ± SEM; n=10-11 experiments). Each data point is the sum of all structures in 20 cells per replicate. (I) Confocal microscopy images of control and VNUT knockdown MIN6 cells immunostained against insulin (green) and proinsulin (magenta). (J) Quantification of the insulin immunoreactivity in control and VNUT KD cells (n=6-8 experiments, 557-677 cells, *P<0.05, **P<0.001, Mann-Whitney U-test). (K) Quantification of the ratio between insulin and proinsulin in control and VNUT knockdown MIN6 cells (means ± SEM; n=5, >60 cells, **P<0.01, Mann-Whitney U- test). (L) Insulin content in control and VNUT KD MIN6 cells determined by ELISA (means ± SEM; n=3). (M) Confocal microscopy images of control and VNUT KD MIN6 cells immunostained against insulin. Images below are magnifications where individual granules are visible. (N) Insulin immunoreactivity in individual insulin granules in control and VNUT KD cells (means±S.D.; N=3 experiments, n= 74 cells per conditions, 20 granules per cell; ***P<0.001, Student’s unpaired t-test). Pictures are averaged images of individual granules in control and VNUT KD cells (200 granules from 20 cells per condition). (O) Proinsulin immunoreactivity in individual insulin granules in control and VNUT KD cells (means±S.E.M.; N=2 experiments, n= 33 cells per conditions, 20 granules per cell, Student’s unpaired t-test).

### VNUT is required for normal insulin secretion

When IGs undergo exocytosis, ATP is co-released together with insulin and act as a local auto- and paracrine regulator through purinergic receptor signaling ^37^. To test if ATP release was impaired in cells with reduced VNUT levels, we expressed mCherry-C1aC1b_PKC_, which reports changes in plasma membrane diacylglycerol (DAG) levels following P2Y1 receptor activation and imaged the cells with TIRF microscopy. As previously shown, KCl-induced depolarization of control cells triggered rapid and transient DAG formation in the plasma membrane (“spikes”), reflecting activation of purinergic receptors by ATP co-secreted with insulin^37^, and this response was reduced in VNUT knockdown cells (Fig. 4A,B). In parallel with the DAG measurements, we also visualized IG fusion with the plasma membrane using the pH-sensitive granule marker vesicle-associated membrane protein 2 (VAMP2)-pHluorin (Fig. 4A). The VAMP2-pHluorin fluorescence is quenched inside the acidic IG lumen. Upon exposure of the vesicle lumen to the pH-neutral extracellular space, the pHluorin fluorescence increases and enable visualization of fusion events. Depolarization of control cells triggered pronounced accumulation of VAMP2-pHluorin in the plasma membrane, reflecting exocytosis, and this response was reduced by around 50% in VNUT knockdown cells (Fig. 4C). To confirm that VNUT knockdown reduced insulin secretion, we employed a real-time optical assay based on feedback activation of insulin receptors with ensuing activation of PI3-kinase and production of plasma membrane PI(3,4,5)P_3_, which was detected with PH_GRP1_-GFP_4_ ^38^ (Fig. 4A). Depolarization triggered robust PI(3,4,5)P_3_ formation in control cells which was reduced, but not absent, in VNUT knockdown cells, consistent with reduced insulin secretion. Measurements of intracellular Ca^2+^ concentration changes using R-GECO1 did not reveal any differences in depolarization-induced Ca^2+^ influx between control and VNUT knockdown cells, showing that the defect in secretion occurs distal to Ca^2+^ influx. (Fig. 4E). Very similar results were obtained when VNUT activity was instead blocked using the inhibitor Clodronate ^39^. Pre- treatment with Clodronate for 12-16 h reduced depolarization-induced IG exocytosis by 50%, but was without effect on the preceding Ca^2+^ influx (Fig. 4F-H). Together, these results show that VNUT activity is required for normal insulin secretion.

### VNUT activity prevents lysosomal degradation of IGs

The reduction of both insulin content and secretion in cells with reduced VNUT expression could be explained by increased IG turnover. To measure lysosomal IG degradation, we developed a live cell reporter where granule-localized NPY was fused to the green fluorescent protein mNeonGreen and the red fluorescent protein mCherry. Inside the IG lumen, where pH is around 6, signal from both fluorophores was readily detected and overlapped. When IGs fuse with lysosomes, where the pH drops to around 5, the mNeonGreen signal was fully quenched while mCherry was still detectable (Fig. S4A). When expressed in mouse islets, the sensor localized both to IGs, seen as small, spherical structures positive for both mNeonGreen and mCherry, and to larger structures that were only positive for mCherry, representing lysosomes (Fig. S4B). When islets were starved for 4 h by removing either glucose or amino acids from the culture medium, this led to an increase in the number and size of mCherry-positive lysosomes, indicating increased IG turnover (Fig. S4B-E). Next, we expressed the reporter in control and VNUT knockdown MIN6 cells and observed them under a confocal microscope. In control cells, most of the reporter signal came from IGs and only few mCherry- positive lysosomes were observed. When VNUT expression was reduced, this was accompanied by a marked increase in both the number and size of mCherry-positive structures as well as an increased mCherry fluorescence intensity inside individual lysosomes, indicating increased IG turnover (Fig. 5A-C). The mCherry-positive structures overlapped with the lysosomal marker SiR-lysosome and were sensitive to the lysosomal inhibitor chloroquine, showing that IG cargo is deposited in lysosomes (Fig. 5D-F). To test if IGs were delivered to lysosomes via autophagy, we treated control and VNUT KD cells for 12-16 h with the Vps34 inhibitor SAR405, which is a potent blocker of macroautophagy ^40^. Consistent with an involvement of autophagy in IG turnover, SAR405 completely prevented the lysosomal accumulation of IG cargo and restored insulin content in VNUT knockdown cells (Fig. 5G-I). Finally, to test if lysosomal degradation of IGs required the ATP transport ability of VNUT, we treated cells with the VNUT inhibitor Clodronate for 2-3 h and determined IG turnover using NPY-mCherry-mNeonGreen. We observed that clodronate increased lysosomal degradation of IGs in control cells but not in VNUT knockdown cells, where turnover was already high (Fig. 5J-L). Together, these results show that loss of VNUT activity direct IGs towards lysosomal degradation.

**Figure 4.**
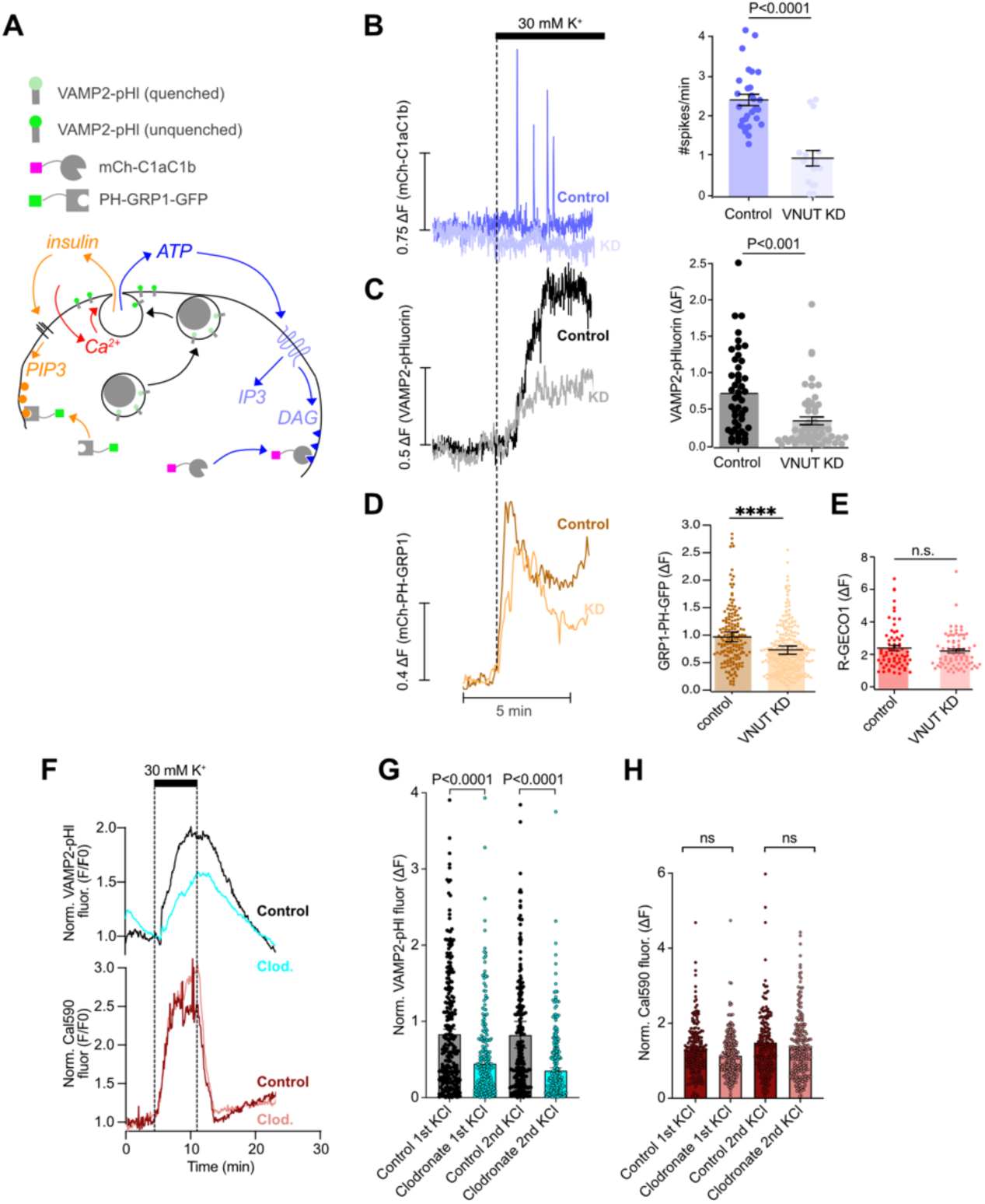
VNUT is required for normal insulin secretion. (A) The principle of detection of fusion events with VAMP2-pHluorin, ATP feedback with mCh-C1aC1b and insulin feedback with PH-GRP1-GFP. (B) TIRF microscopy recordings of mCh-C1aC1b fluorescence change in response to a brief depolarization in control (dark) and VNUT knockdown (light) MIN6 cells (control: n=28 cells, 3 replicates; VNUT KD: n=16 cells, 3 replicates; Student’s 2-tailed unpaired t-test). (C) TIRF microscopy recordings of VAMP2-pHluorin fluorescence change in response to a brief depolarization in control (dark) and VNUT knockdown (light) MIN6 cells (control: n=48 cells, 5 replicates; VNUT KD: n=52 cells, 5 replicates; Student’s 2-tailed unpaired t-test). (D) TIRF microscopy recordings of GRP1-PH-GFP fluorescence change in response to depolarization in control (dark) and VNUT knockdown (light) MIN6 cells (n=86-117 cells from three experiments; ****P<0.001, Student’s unpaired t-test). (E) R-GECO1 fluorescence increase in response to depolarization in control (dark) and VNUT knockdown (light) MIN6 cells (n=67-71 cells from two experiments; Student’s unpaired t-test). (F) TIRF microscopy recordings of VAMP2-pHluorin (blue) and Cal590 (red) fluorescence change in response to depolarization in control MIN6 cells (dark) and cells treated for 2-3 h with 120 nM clodronate (light). (G) Quantification of VAMP2-pHluorin fluorescence increase at the plasma membrane following two brief depolarizations 20 min apart (KCl1 and KCl2) in control MIN6 cells (dark) and cells treated for 2-3 h with 120 nM clodronate (light) (n=242-290 cells from 5 experiments, Student’s unpaired t-test). (H) Quantification of Cal590 fluorescence increase following two brief depolarizations 20 min apart (KCl1 and KCl2) in control MIN6 cells (dark) and cells treated for 2-3 h with 120 nM clodronate (light) (n=202-235 cells from 3 experiments, Student’s unpaired t- test).

**Figure 5.**
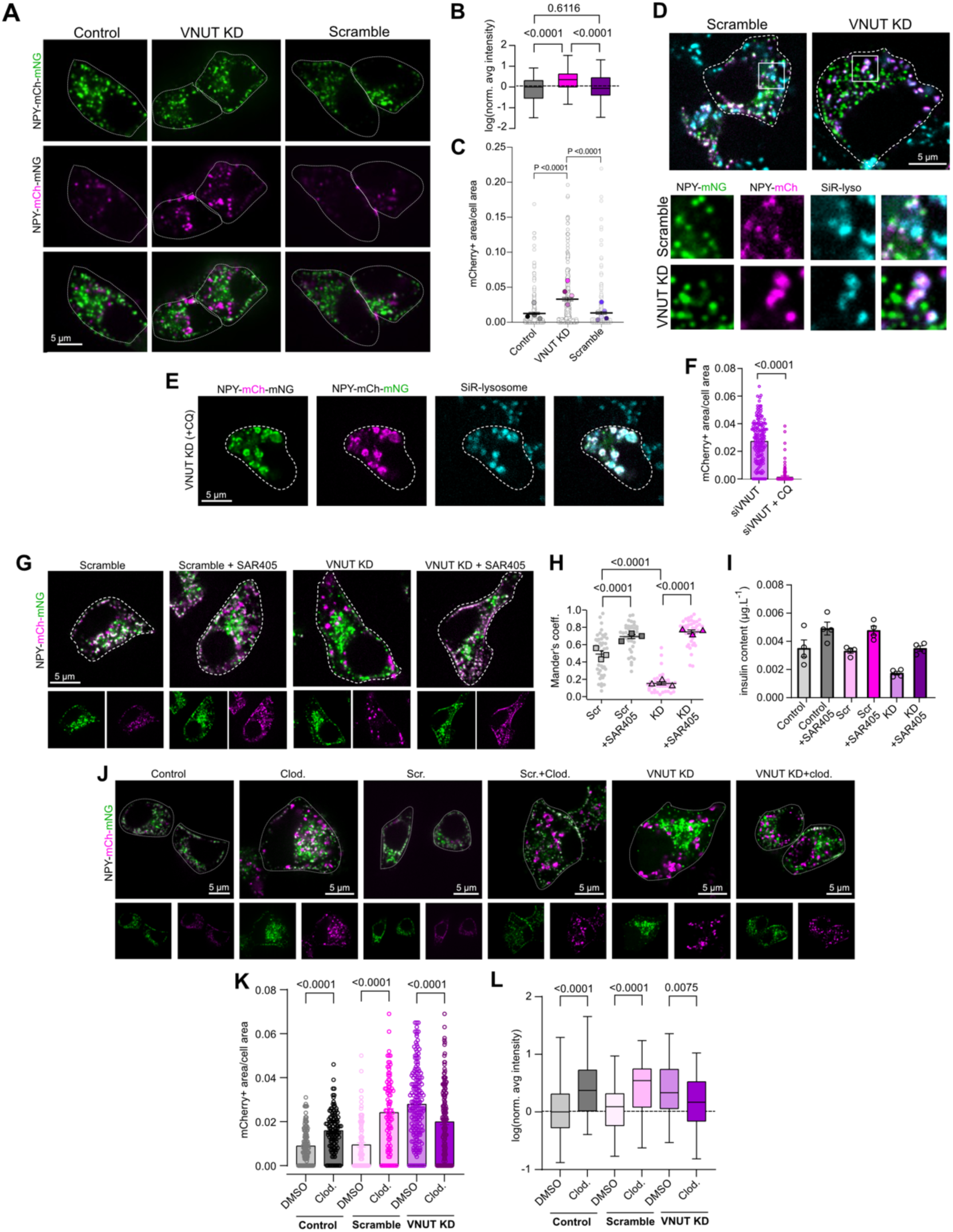
VNUT knockdown results in lysosomal degradation of insulin granules. (A) Confocal microscopy images of control MIN6 cells or MIN6 cells treated with VNUT or control (scramble) siRNA. Cells are expressing an insulin granule-localized autophagy reporter. (B) Fluoresence intensity of mCherry-positive structures in control and VNUT KD MIN6 cells (line represent average, box is 25th-75^th^ percentiles and whiskers show min-max) (n = 286-371 from 3 experiments, Kruskal-Wallis test -Dunn’s multiple comaprision). (C) mCherry-positive area in control and VNUT KD MIN6 cells (n=263-286 cells from four experiments, Mann-Whitney U-test). (D) Confocal microscopy images of control and VNUT KD MIN6 cells expressing an insulin granule-localized autophagy reporter. Lysosomes are labelled with SiR-lysosome (blue). (E) Confocal microscopy images of a VNUT KD MIN6 cell expressing an insulin granule- localized autophagy reporter following 12-16 h treatment with 20 μM chloroquine. Lysosomes are labelled with SiR-lysosome (blue). (F) mCherry-positive area in VNUT KD MIN6 cells treated with chloroquine (n = 146-170 from 3 experiments, Mann-Whitney U-test) (G) Confocal microscopy images of control and VNUT KD MIN6 cells expressing an insulin granule-localized autophagy reporter in the absence or presence of 10 μM SAR405 (12-16 h treatment). (H) Mander’s overlap coefficient for mCherry and mNeonGreen in control and VNUT KD MIN6 cells expressing NPY-mCherry-mNeonGreen and treated with SAR405 (n=3 experiments, 42 cells, Mann-Whitney U-test). (I) Insulin content in control and VNUT KD MIN6 cells treated or not with 10 µM SAR405 for 12-16 h (n=4). (J) Confocal microscopy images of control and VNUT KD MIN6 cells treated or not with 120 nM clodronate for 2-3 h. (K) mCherry-positive area in control and VNUT KD MIN6 cells treated or not with 120 nM clodronate (n = 91-173 from 3 experiments, Mann-Whitney U-test). (L) Fluorescence intensity of mCherry-positive structures in control and VNUT KD MIN6 cells treated or not with 120 nM clodronate (line represent average, box is 25th-75^th^ percentiles and whiskers show min-max; n = 91-173 from 3 experiments, Mann- Whitney U-test).

## DISCUSSION

Beta cells have the ability to match the demands of insulin release with the production of IGs^21^ , and although beta cells only secrete around 5% of the total insulin content per hour ^41,42^, most of this insulin is stored in granules that has been recently produced, highlighting the importance of a functional insulin biosynthetic pathway ^13,43^. Maturation of newly formed IGs involves changes in both protein and lipid composition, as well as acidification of the granule lumen, accumulation of Ca^2+^, ATP and other signaling molecules, and processing of proinsulin to insulin ^8,10,44,45^. All these events occur within an hour after granule budding from the TGN, indicating that it is a highly regulated and coordinated process ^13^. Lipid, ionic and metabolite uptake are known to be concentrated to membrane contact sites, where transfer between two organelles is facilitated by their close proximity ^46^. We recently found that IGs form contacts with the ER as part of the maturation process ^15^, and in the current work we now show that IGs also directly communicate with mitochondria. The putative existence of IG- mitochondria complexes was first shown almost 50 years ago ^20^, and more recent work has confirmed their existence ^19,34^, but nothing is known about the composition or function of these contacts. Using a combination of fixed and live-cell proximity detection techniques we found that recently synthesized IGs, located close to the trans-Golgi network, interact with mitochondria, and that this interaction likely involves granular VNUT and mitochondrial VDAC1. While VDAC1 is involved in tethering mitochondria to numerous organelles, including the ER^46^, solute carriers are not typically part of organelle tethers, although they can be enriched at MCS ^47^. VDAC is a general carrier that forms a pore in the outer mitochondrial membrane that permits passage of metabolites, including ATP, and coupling this transport to ATP uptake through granular VNUT may greatly facilitate this critical step in granule maturation. ATP produced by granule-proximal mitochondria may also facilitate energy- dependent processes, such as clathrin uncoating, maintenance of Ca^2+^ and pH gradients, or uptake of other metabolites, as has been shown for fatty acid uptake into the ER, which depends on ATP locally generated at ER-mitochondria contacts ^48^. Additional functions of these contacts beyond local ATP delivery are also likely. A recent study identified two populations of IGs characterized by high and low reactive oxygen species (ROS) content, where the more oxidized pool localized at the cell center, resembling the distribution of IG- mitochondria contacts ^49^. Given that mitochondria are the main source of ROS, it is tempting to speculate that ROS transfer to granules may occur at mitochondrial contact sites, similar to what has been shown for peroxisomes, which are the main ROS-scavenging organelle in non- secretory cells ^50^.

ATP is a powerful hydrotrop and an osmotic agent that help to concentrate hormones, and VNUT-mediated ATP uptake into secretory granules is therefore critical for packaging of cargo proteins ^36,51^. Consistent with previous studies ^36,52,53^, we show that reduced expression or pharmacological inhibition of VNUT leads to reduced granular ATP content and to reduced hormone secretion ^36,52^. The effect on hormone secretion has been ascribed to both impaired autocrine feedback ^52^ and reduced number of granule fusion events at the plasma membrane^36^. Here, we observe a marked reduction in insulin content in cells with reduced VNUT expression, and time-dependent labelling of IGs showed a preferential loss of recently synthesized granules over older ones. VNUT has been shown to localize to a subset of IGs defined by the presence of the high-affinity Ca^2+^ sensor synaptotagmin-7 and high cholesterol content ^54^. These granules have been proposed to be recruited to release sites during sustained insulin secretion, and likely correspond to granules that have recently formed at the TGN ^8^. It is therefore possible that granules that retain VNUT following budding from the TGN, and subsequently interact with mitochondria and accumulate ATP, are specifically directed towards exocytosis, and that loss of VNUT eliminate this granule pool, leading to impaired insulin secretion. Since insulin content was dramatically reduced in cells with reduced VNUT expression, we suspected that granules were lost through lysosomal degradation. It is well- known that IGs that fail to undergo exocytosis are eventually degraded in lysosomes ^55,56^, and that the probability increases with granule age ^9^, but the triggering signal for degradation is not known. To follow IG turnover, we adapted an approach previously used to monitor mitophagy ^57^, and found that loss of VNUT resulted in deposition of IG cargo into lysosomes, resulting in both more granule cargo inside lysosomes and increased number of lysosomes containing granule cargo. Suppression of macroautophagy with the VPS34 inhibitor SAR405 ^40^ completely prevented the lysosomal accumulation of granule cargo in VNUT deficient cells, indicating that enhanced autophagy is the cause of reduced insulin content in these cells. Chlodronate, a pharmacological inhibitor of VNUT ^39^, also strongly increased lysosomal degradation of IGs, showing that it is the ATP transport activity, and not putative mitochondria tethering function, of VNUT that is required for normal progression through the secretory pathway.

Diabetes and diabetes-like conditions increase autophagic removal of IGs^22^, while at the same time compromising mitochondrial function, preventing both mitochondrial interactions with other organelles as well as reducing ATP production^17,58^. In addition to the acute negative effects of ATP deficiency on insulin secretion, it is likely that these pathological changes will prevent mitochondria-IG crosstalk and lead to reduced granular ATP content, thus contributing to lysosomal degradation of IGs and in insulin deficiency. While numerous mechanisms that promote lysosomal degradation of newly formed insulin granules has been described ^22,59,60^, their relative contribution and dependence on each other is unclear. Perhaps ATP-loss is a hallmark of aging or dysfunctional granules that act as a quality control mechanism to determine if a granule should be directed towards degradation or secretion. To test this hypothesis will be an important future research goal. Additionally, it will be important to more carefully characterize the mitochondria-IG contacts and to determine their importance for both IG biogenesis and mitochondrial function. While not a main aim in the present study, our results show that both mitochondrial morphology and membrane potential was affected by the loss of granule-localized VNUT, indicating that granules may control mitochondrial function and not only vice versa.

