## Supplementary figures for "Mitochondria – insulin granule crosstalk controls the early stages of granule maturation"

**SUPPLEMENTARY FIGURES AND8 LEGENDS**

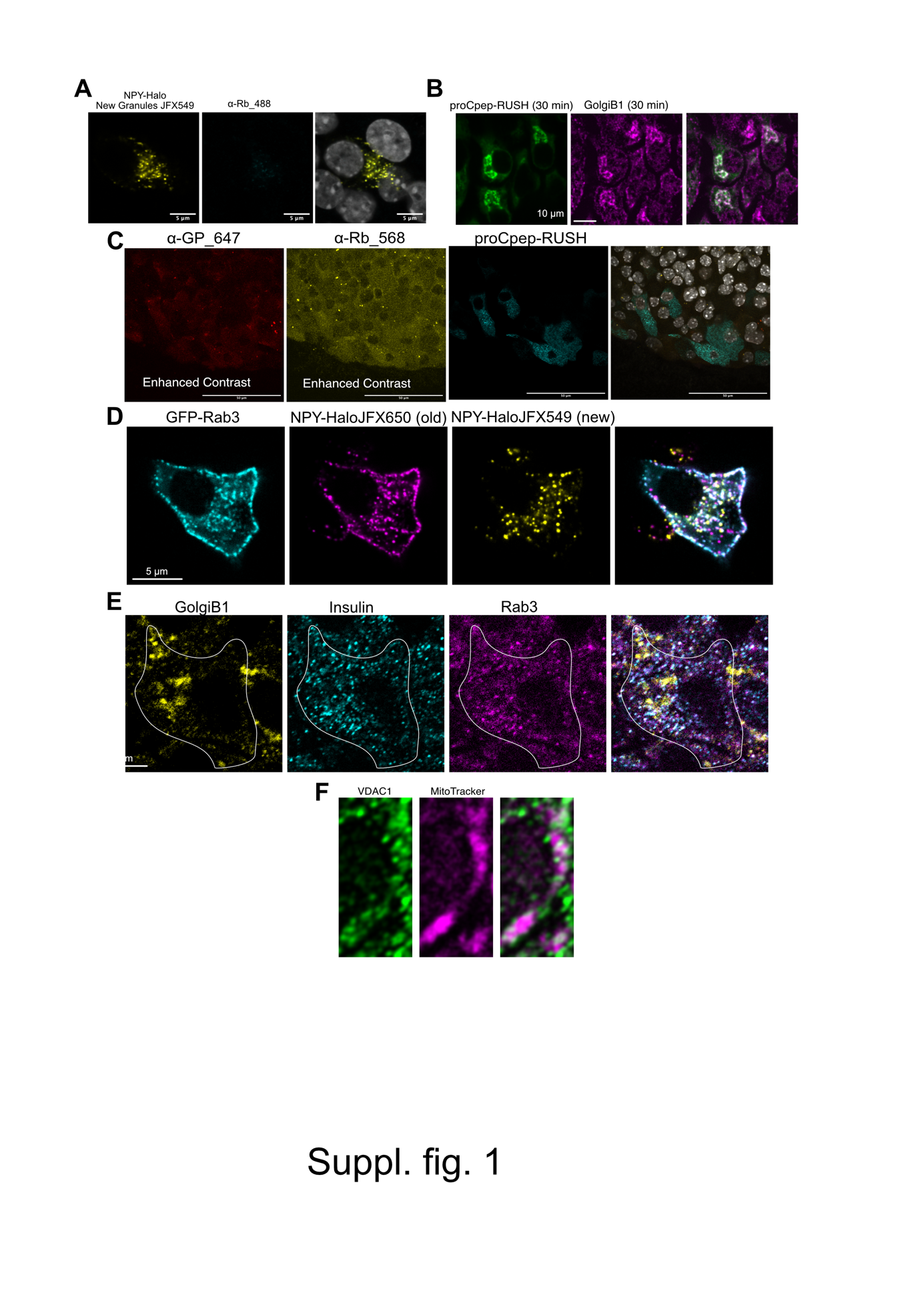

**Suppl. Fig. 1.**

(A) Confocal microscopy image of MIN6 cells expressing NPY-Halo labelled with JFX549, fixed, permeabilized and exposed to α-Rabbit alexa-488 secondary antibody. Notice the absence of non-specific background signal from the secondary antibody.
(B) Confocal miroscopy images of MIN6 cells expressing proCpep-RUSH following 30 min incubation with biotin followed by fixation and immunolabeling of the Golgi with a GolgiB1 antibody.
(C) Confocal microscopy images of fixed and permeabilized mouse islets expressing proCpep-RUSH and exposed to the indicated secondary antibodies (α-Guinea pig alexa-647 and α-Rabbit alexa-568). Notice the absence of specific immunoreactivity from the secondary antibodies (brightness/contrast has been adjusted to clearly show the weak, non-specific signal from the secondary antibodies).
(D) Confocal microscopy images of a MIN6 cell expressing GFP-Rab3 (cyan) and NPY-Halo that has been pulse-chase labeled with JFX650 (magenta; old granules) and JFX549 (yellow; new granules).
(E) Confocal microscopy images of MIN6 cells immunostained against GolgiB1 (yellow), insulin (cyan) and Rab3 (magenta).
(F) Confocal microscopy image of subcellular region of a MIN6 cell showing the distribution of VDAC1 (green) in relation to the mitochondrial dye MitoTracker deep red.

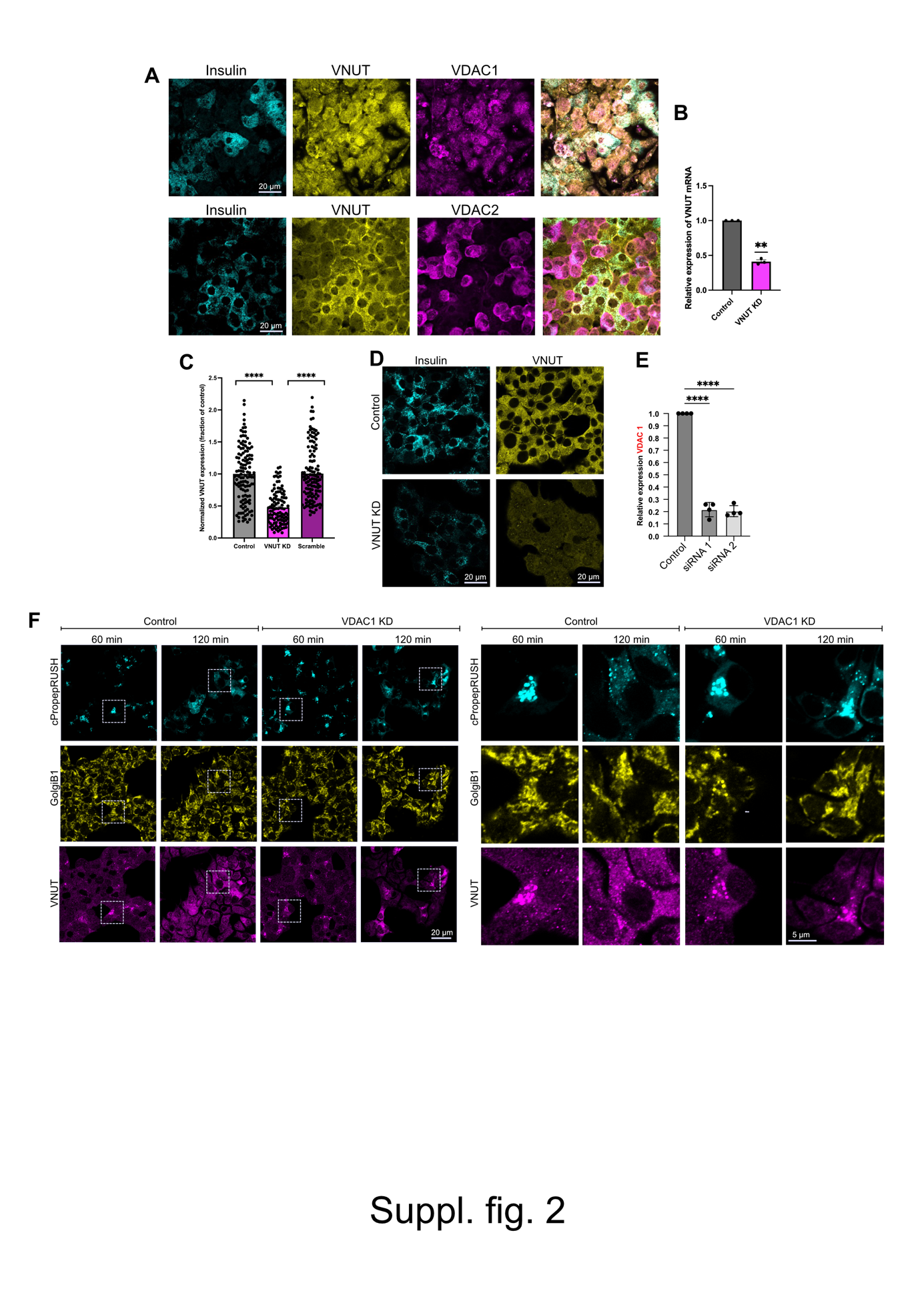

**Suppl. Fig. 2**

(A) Confocal microscopy images of mouse islets immunostained against insulin (cyan), VNUT (yellow) and VDAC1 (magenta; top) or VDAC2 (magenta; bottom).
(B) Quantitative RT-PCR determination of VNUT mRNA levels in control (grey) and VNUT knockdown cells (magenta, expressed relative to control after normalization to GAPDH mRNA levels (n=3, ** P<0.01).
(C) Quantification of insulin immunoreactivity in control MIN6 cells and MIN6 cells treated with control (scramble) or VNUT siRNA. Data is expressed relative to control (n=150-160 cells from three experiments; ****P<0.001, Student’s unpaired t-test).
(D) Confocal microscopy images of control and VNUT KD MIN6 cells immunostained against insulin (cyan) and VNUT (yellow).
(E) Quantitative RT-PCR determination of VDAC1 mRNA levels in control and VDAC1 knockdown cells. Data is expressed relative to control after normalization to GAPDH mRNA levels (n=4, **** P<0.001).
(F) Confocal microscopy images of control and VDAC1 KD MIN6 cells expressing cPropep-RUSH that has been treated with biotin for 60 or 120 min, followed by fixation and immunostaining against GolgiB1 (yellow) and VNUT (magenta). The boxed region is magnified to the right.

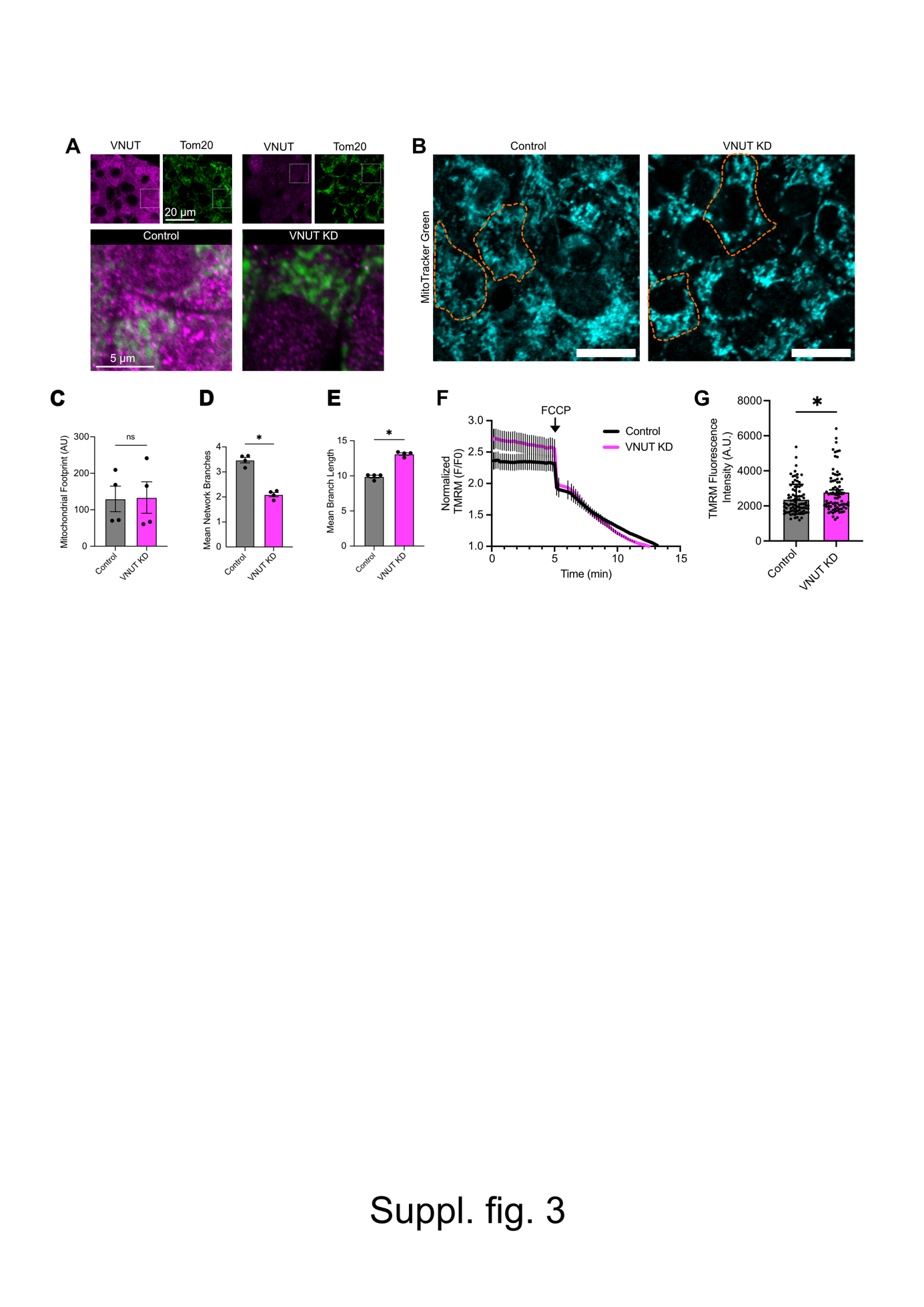

**Suppl. Fig. 3**

(A) Confocal microscopy images of control and VNUT KD MIN6 cells immunostained against VNUT (magenta) and Tom20 (green; mitochondria). Images below are magnifications.
(B) Confocal microscopy images of control and VNUT knockdown MIN6 cells loaded with Mitotracker-green. The orange dashed lines highlight two cells. Scale bar is 10 µm.
(C) The mitochondrial footprint in control and VNUT knockdown MIN6 cells (mean±S.E.M.; n=4).
(D) Number of mitochondrial branches in control and VNUT knockdown MIN6 cells (mean±S.E.M.; n=4; *P<0.05).
(E) Mitochondrial branch length in control and VNUT knockdown MIN6 cells (mean±S.E.M.; n=4; *P<0.05).
(F) TMRM fluorescence change in control (black) and VNUT knockdown (magenta) MIN6 cells kept in 20 mM glucose and 250 µM diazoxide and exposed to 1 µM FCCP. The signal is normalized to the lowest value obtained in the presence of FCCP (means ± SD; 70-80 cells, ***P<0.01).
(G) Absolute TMRM fluorescence in control and VNUT knockdown MIN6 cells kept in a buffer containing 20 mM glucose and 250 µM diazoxide (means ± SD; 70-80 cells, ***P<0.01).

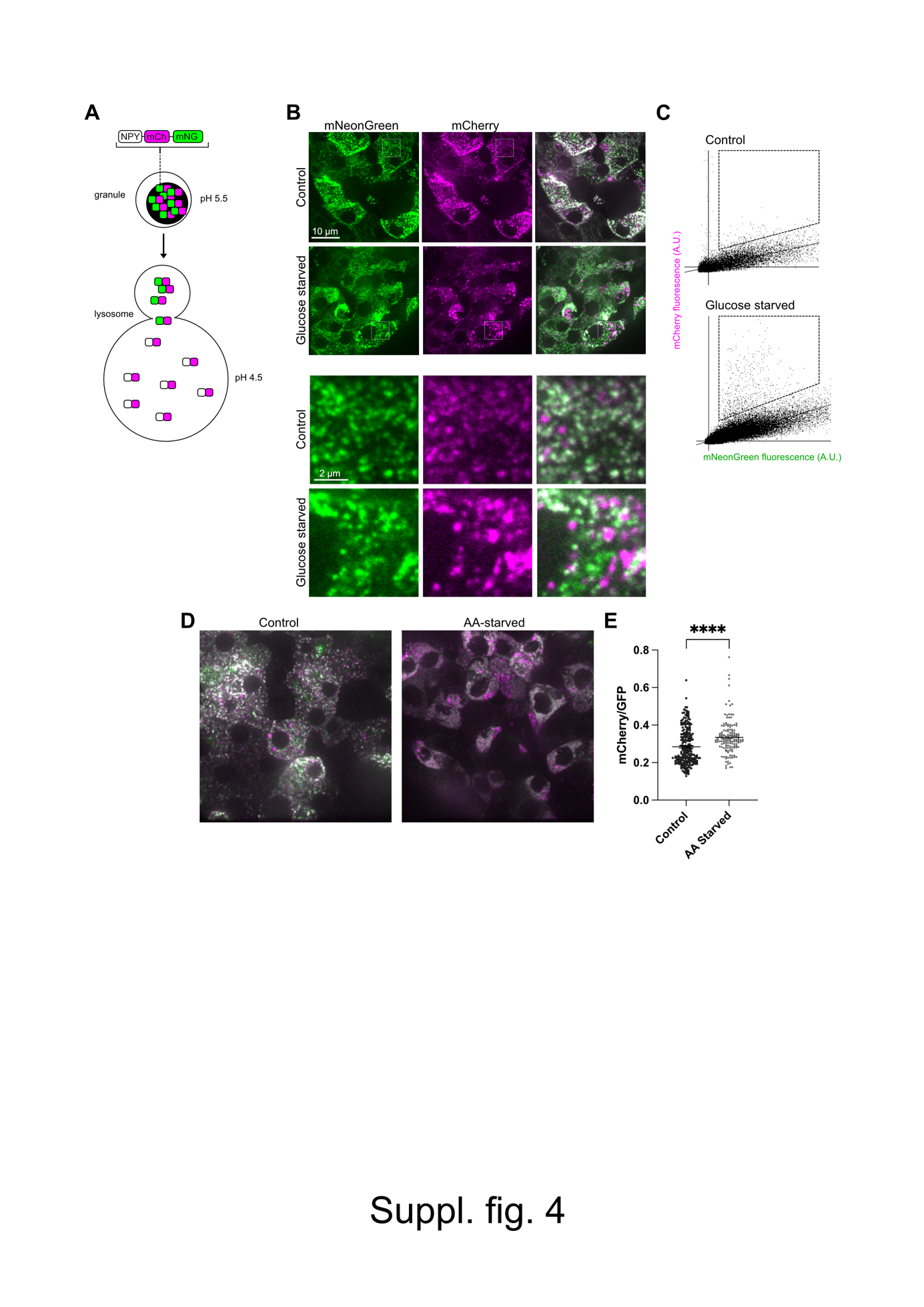

**Suppl. Fig. 4**

(A) A fluorescent biosensor to detect lysosomal degradation of insulin granules. The biosensor is composed of NPY fused to pH-sensitive mNeonGreen and pH-insensitive mCherry. Both fluorophores are fluorescent in the granule lumen, but only mCherry retain this propertiy inside the lumen of the highly acidic lysosme. Therefore, lysosomes containing granule cargo can be detected with fluorescence microscopy as mCherry-positive, mNeonGreen-negative objects.
(B) Confocal microscopy images of mouse islets expressing NPY-mNG-mCherry. The islets have been cultured in normal medium with 10 mM glucose (control) or in a glucose-deficient medium (glucose starved) for 4 h prior to imaging. The content of the dashed white boxes are magnified below. Notice the appearence of large, mCherry-positive, mNG-negative structures in the glucose-starved islet cells.
(C) Scatter plot showing pixel intesity distributions in the islet in (B). Notice an increase in pixel numbers inside the boxed area that corresponds to mCherry+/mNG- pixels in the glucose-starved islet.
(D) Confocal microscopy images of mouse islets expressing NPY-mNG-mCherry. The islets have been cultured in normal medium with 2 mM L-glutamine (control) or in a amino acid-deficient medium (AA-starved) for 4 h prior to imaging. Notice the appearence of large, mCherry-positive, mNG-negative structures in the AA-starved islet cells.
(E) mCherry/GFP fluorescence ratio in cells from islets cultured under normal conditions (control) or in the absence of amino acids (AA-starved) (n=>100 cells from two islet preparations; ****P<0.001; Mann-Whitney U-test).
